# An RNA replicon vaccine encoding HA and NA prevents shedding of antigen-drifted 2009 pandemic H1N1 influenza virus in the pig model

**DOI:** 10.1101/2025.11.01.685547

**Authors:** Obdulio Garcia-Nicolas, Lisa Butticaz, Robin Avanthay, Nicolas Ruggli, Artur Summerfield, Gert Zimmer

## Abstract

Seasonal influenza viruses escape the human immune response by antigenic drift, i.e. the positive selection of point mutations that prevent the binding of inhibitory antibodies to the influenza antigens HA and NA. The efficacy of seasonal influenza vaccines can be less than 50% if the selected influenza vaccine strain does not match the antigenic characteristics of the circulating seasonal influenza virus. In this study, we used the porcine model to evaluate the efficacy of an RNA replicon vaccine encoding the HA and NA antigens of A/Hamburg/4/2009 (H1N1) (H1N1_HH4/09_) in inducing cross-reactive immunity. We found that a single intramuscular immunization with this vaccine elicited high levels of antibodies with H1N1_HH4/09_-neutralizing activity and potent N1-sialidase inhibition. A second immunization with the same H1/N1 RNA replicon particles or with a live-attenuated influenza vaccine (LAIV) based on a modified H1N1_HH4/09_ virus boosted the inhibitory activity of the immune sera against the antigen-drifted A/Victoria/2570/2019 (H1N1) (H1N1_Vic/19_) strain. Interestingly, vaccination elicited N1-specific antibodies that also inhibited the activity of avian N1 sialidase and potently inhibited the replication of A/cattle/Texas/063224-24-1/2024 (H5N1) (H5N1_Tex/24_) *in vitro*. When challenged nasally with a H1N1_HH4/09_ /H1N1_Vic/19_ 6:2 reassortant virus encoding the HA and NA antigens of H1N1_Vic/19_, immunized pigs did not shed infectious virus while the control animals did, suggesting that homologous prime/boost vaccination with H1/N1 replicon particles can block virus replication in the upper respiratory tract as efficiently as the heterologous RNA replicon prime/LAIV boost immunization regimen. In conclusion, RNA replicons encoding both HA and NA either used alone or in combination with LAIV mediate protection against antigen-drifted influenza viruses and reduce the risk of vaccination breakthroughs due to antigen mismatch. Furthermore, this vaccine may also limit the infection by zoonotic H5N1 viruses.

## Introduction

Influenza A viruses (IAV) encode two important envelope glycoproteins that are targeted by the immune system of the host. The hemagglutinin (HA) is a trimeric transmembrane protein with a large ectodomain that is structured into a variable globular head domain and a more conserved stalk region. The globular head domain harbors the receptor-binding pocket that interacts with sialic acid residues on cellular glycoproteins and glycolipids ^1^. Most virus-neutralizing antibodies that are produced by the immune system in response to IAV infection are directed to the globular head and interfere with the receptor-binding activity of HA ^1,2^. However, IAV are highly mutable, and the immune response frequently selects for viral mutants that are characterized by amino acid substitutions in the globular head region, which prevents binding of neutralizing antibodies, a phenomenon known as “antigenic drift” ^3^. Because of this antigenic drift, seasonal influenza vaccines need to be updated every year according to WHO recommendations which are published about six months before the upcoming influenza season. Sometimes this prediction is inaccurate and a mismatch between the vaccine strain and presently circulating seasonal IAV leads to low vaccine efficacy ^4,5^. Thus, there is a need for vaccines that provide efficient protection against antigen-drifted IAV.

Neuraminidase (NA) is the second important antigen of IAV. The tetrameric transmembrane protein exhibits sialidase activity which is essentially required for release of progeny virus from the infected cell membrane ^6^, as well as for penetration of the mucus-rich layers covering the respiratory epithelia of the host ^7^. Antibodies that inhibit NA sialidase activity can efficiently interfere with virus dissemination and thus have protective potential ^8–10^. Not surprisingly, NA is also subject to antigenic drift ^11–13^. Of note, the most used vaccines to control seasonal influenza epidemics are inactivated split IAV standardized for hemagglutinin (HA) content, whereas the protective features of the NA are usually neglected ^14^.

In our previous work, we developed a novel prime/boost vaccination strategy comprising of an intramuscular prime with recombinant RNA replicon particles composed of G protein-deleted vesicular stomatitis virus (VSVΔG) vector encoding the H1 antigen of the pandemic A/Hamburg/4/2009 (H1N1) strain (H1N1_HH4/09_) followed by an intranasal boost using a live-attenuated influenza vaccine (LAIV) based on a modified H1N1_HH4/09_ characterized by a truncated NS1 protein and lack of PA-X protein expression (NS1(1-126)-ΔPAX) ^15^. This prime/boost vaccination protocol elicited a strong systemic and mucosal immune response in the porcine model and resulted in sterilizing immunity against challenge infection with the homotypic H1N1_HH4/09_ strain. In addition, priming with H1-recombinant RNA replicon particles controls replication and shedding of LAIV, thereby adding to vaccine safety ^16^.

In the present study, we investigated the efficacy of the heterologous RNA replicon prime/LAIV boost strategy in protecting pigs against antigen-drifted H1N1 virus and compared it with a homologous RNA replicon prime/RNA replicon boost strategy. To achieve a broader immune response, a cocktail of two VSV-based RNA replicon particles was used for intramuscular priming of the animals, VSVΔG(H1) and VSVΔG(N1), which encode the HA and NA antigens of H1N1_HH4/09_, respectively. In the homologous prime/boost strategy, the animals were immunized a second time with the RNA replicon particles, while in the heterologous strategy, they were immunized intranasally with LAIV NS1(1-126)-ΔPAX. The immune response of the animals was examined for H1-specific virus-neutralizing antibodies as well as N1-specific antibodies showing sialidase-inhibitory activity. Finally, the animals were challenged with a reassortant virus encoding the HA and NA antigens of the antigen-drifted A/Victoria/2570/2019 (H1N1) (H1N1_Vic/19_) strain. Our results suggest that while both prime/boost strategies efficiently controlled replication and nasal shedding of the antigen-drifted virus, the robust N1-specific response induced by the VSV replicons turned out to be broadly protective, also inhibiting zoonotic H5N1 virus isolated from infected dairy cows.

## Results

### Evaluation of homologous and heterologous prime/boost vaccination strategies in the pig animal model

Specific pathogen-free (SPF) pigs (8 to 10 weeks of age) were randomly distributed into 4 groups with 5 animals each (**Fig. 1**). Animals of groups A and C were first immunized via the intramuscular (i.m.) route with VSV*ΔG, RNA replicon particles encoding the irrelevant GFP protein, while animals of groups C and D were immunized (i.m.) with VSVΔG(H1) and VSVΔG(N1) replicon particles (**Fig. 1**). Four weeks after the primary immunization, the pigs were boosted with either VSV*ΔG (group A) or VSVΔG(H1) and VSVΔG(N1) replicon particles (group B), while animals of groups C and D were boosted via the intranasal route with the LAIV NS1(1-126)-ΔPAX ^15,16^. Vaccination of the animals with VSV replicon particles or LAIV was well tolerated by the animals and did not cause any significant clinical symptoms (**Supplementary Fig. 1**, **Supplementary Tab. 1**). Nasal swab samples were collected on the following days from the animals of groups C and D to monitor shedding of the LAIV. In addition, three immunologically naïve pigs were added to each group C and group D two days after the booster vaccination to monitor potential transmission of the virus to the sentinel animals (**Fig. 1**).

**Fig. 1.**
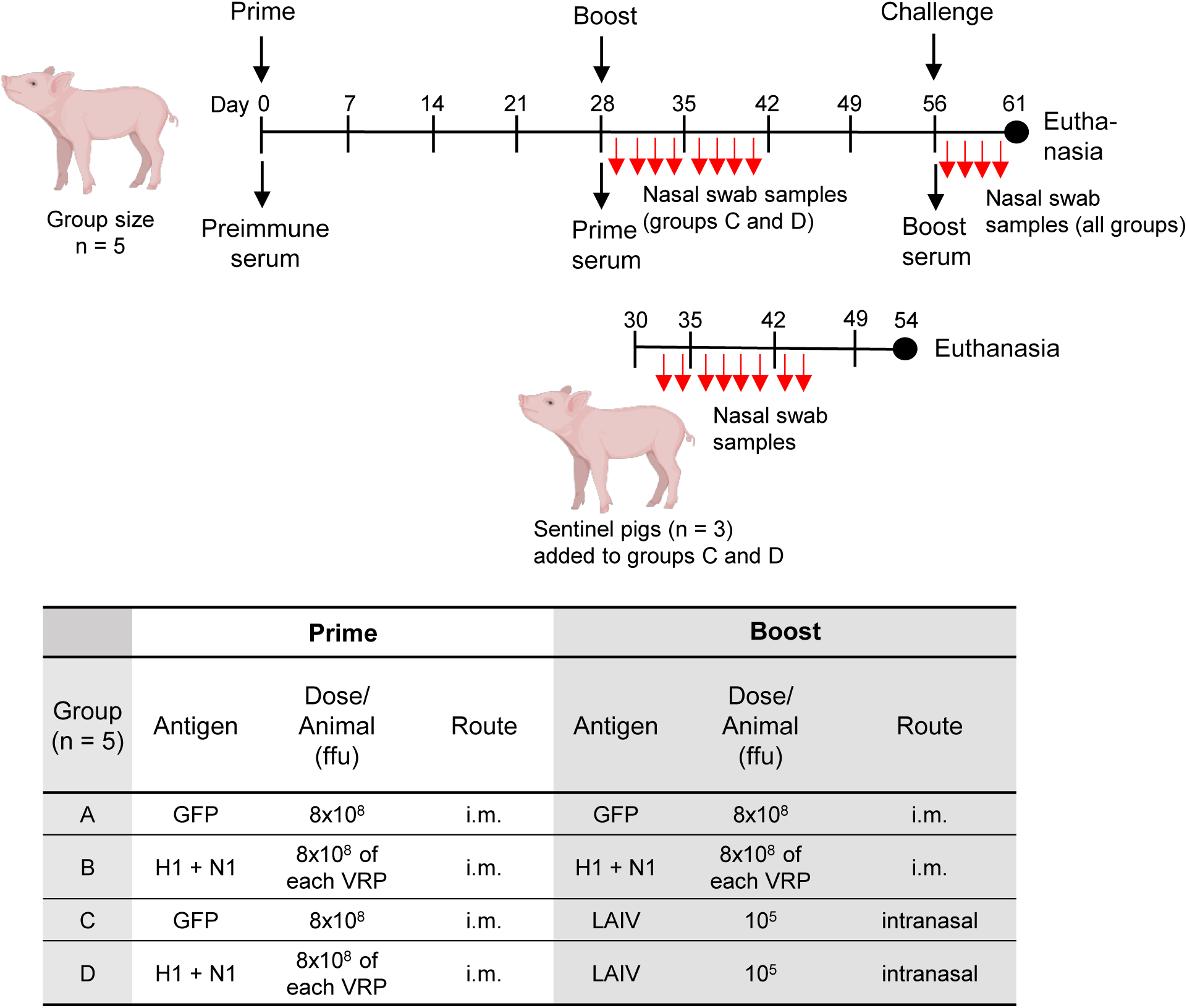
Experimental setup of the prime/boost vaccination regimens used in this study. SPF pigs (8-weeks-old) of mixed sex were randomly assigned to the indicated vaccine groups (n = 5 pigs/group) and were first immunized (i.m.) with either VSV*ΔG (groups A and C) or with VSVΔG(H1) and VSVΔG(N1) replicon particles (groups B and D) using the indicated doses. Four weeks after the primary immunization, the animals were boosted with either the virus-vectored H1 and N1 antigens (i.m.) or the LAIV (intranasal). Sentinel pigs (n = 3) were co-housed on day 30 with groups C and D to monitor transmission of LAIV. Four weeks after the booster vaccination, all animals of groups A to D were challenged intranasally with the H1N1_HH4/09_/H1N1_Vic/19_ 6:2 reassortant virus. Five days post challenge, all animals were euthanized, and lung biopsy samples were taken to determine viral RNA load by RT-qPCR. At the indicated times, blood samples were collected and sera prepared which were analyzed for virus-neutralizing and sialidase-inhibitory activity. In addition, oropharyngeal swab samples were collected (red arrows) and analyzed for the presence of infectious virus and viral RNA load. The piglet image was created in BioRender. Zimmer, G. (2025) https://BioRender.com/ai29p9b.

Four weeks after the second vaccination, the pigs were infected via the intranasal route with a reassortant virus containing the genomic segments 4 (HA) and 6 (NA) of A/Victoria/2570/2019 (H1N1) (H1N1_Vic/19_) and the remaining 6 genomic segments of A/Hamburg/4/2009 (H1N1) (H1N1_HH4/09_). The Victoria strain has been recommended as a component of the tetravalent influenza vaccine for use in the 2021 and 2022 southern hemisphere seasons ^17^ and for the 2022/2023 season in the northern hemisphere ^18^. Compared to the original 2009 pandemic H1N1 virus, the HA and NA antigens of this 2019 H1N1 virus were characterized by several amino acid substitutions in their ectodomains (**Supplementary Figs. 2, 3**).

### The second immunization boosts the neutralizing antibody response against antigen-drifted H1N1 virus

To monitor H1N1-specific antibody responses, serum samples were prepared prior to vaccination (d0, pre-immune serum), 4 weeks after the primary vaccination (d28, prime serum) and 4 weeks after the second immunization (d56, boost serum) (**Fig. 1**), and analyzed for virus-neutralizing activity against H1N1_HH4/09_ (**Fig. 2a**) and the H1N1_HH4/09_/H1N1_Vic/19_ 6:2 reassortant virus encoding the segments 4 and 6 of H1N1_Vic/19_ (**Fig. 2b**). None of the SPF pigs had pre-existing neutralizing antibodies to either of the two viruses. Pigs of group A that had been immunized with the control vaccine VSV*ΔG did not develop any H1N1-neutralizing antibodies, as expected (**Fig. 2a, b**). In contrast, animals of group B developed significantly high titers of H1N1_HH4/09_-neutralizing antibodies (geometric mean ND_50_ = 485) after the first immunization with VSVΔG(H1) and VSVΔG(N1) replicon particles. The ND_50_ titer increased further following the second immunization with the same vaccine (geom. mean ND_50_ titer = 1358), however, the difference between the ND_50_ titers of the prime and boost sera was not statistically significant (**Fig. 2a**). When the immune sera of group B were tested for neutralizing activity against the H1N1_HH4/09_/H1N1_Vic/19_ 6:2 reassortant virus, the prime sera showed only very low neutralizing activity (geom. mean ND_50_ = 6), which increased significantly after the second administration of the vaccine (geom. mean ND_50_ = 277) (**Fig. 2b**). When pigs were first immunized (i.m.) with VSV*ΔG and subsequently with the LAIV via the intranasal route (group C, d56), only low H1N1_HH4/09_-neutralizing serum antibody titers were detected (geom. mean ND_50_ = 30) (**Fig. 2a**), and inhibitory activity against the H1N1_HH4/09_/H1N1_Vic/19_ 6:2 reassortant virus was even lower (geom. mean ND_50_ = 4) (**Fig. 2b**). However, when the animals were first primed (i.m.) with VSVΔG(H1) and VSVΔG(N1) and then intranasally immunized with the LAIV (group D), geometric mean ND_50_ values against the antigen-drifted H1N1_HH4/09_/H1N1_Vic/19_ 6:2 reassortant virus increased from 6 (d28, after prime) to 276 (d56, after boost) (**Fig. 2b**), while neutralizing activity against the homotypic H1N1_HH4/09_ strain was already high after the first immunization (geom. mean ND_50_ = 365), and increased only marginally following immunization with LAIV (geom. mean ND_50_ = 958) (**Fig. 2a**). Finally, we tested whether the boost sera of group B would show some neutralizing activity against A/cattle/Texas/063224-24-1/2024 (H5N1) (H5N1_Tex/24_), however, no significant inhibition of this virus was observed (**Fig. 2c**). In conclusion, our findings suggest that a prime/boost vaccination regimen is important to produce robust levels of serum antibodies with neutralizing activity against antigen-drifted H1N1 virus.

**Fig. 2.**
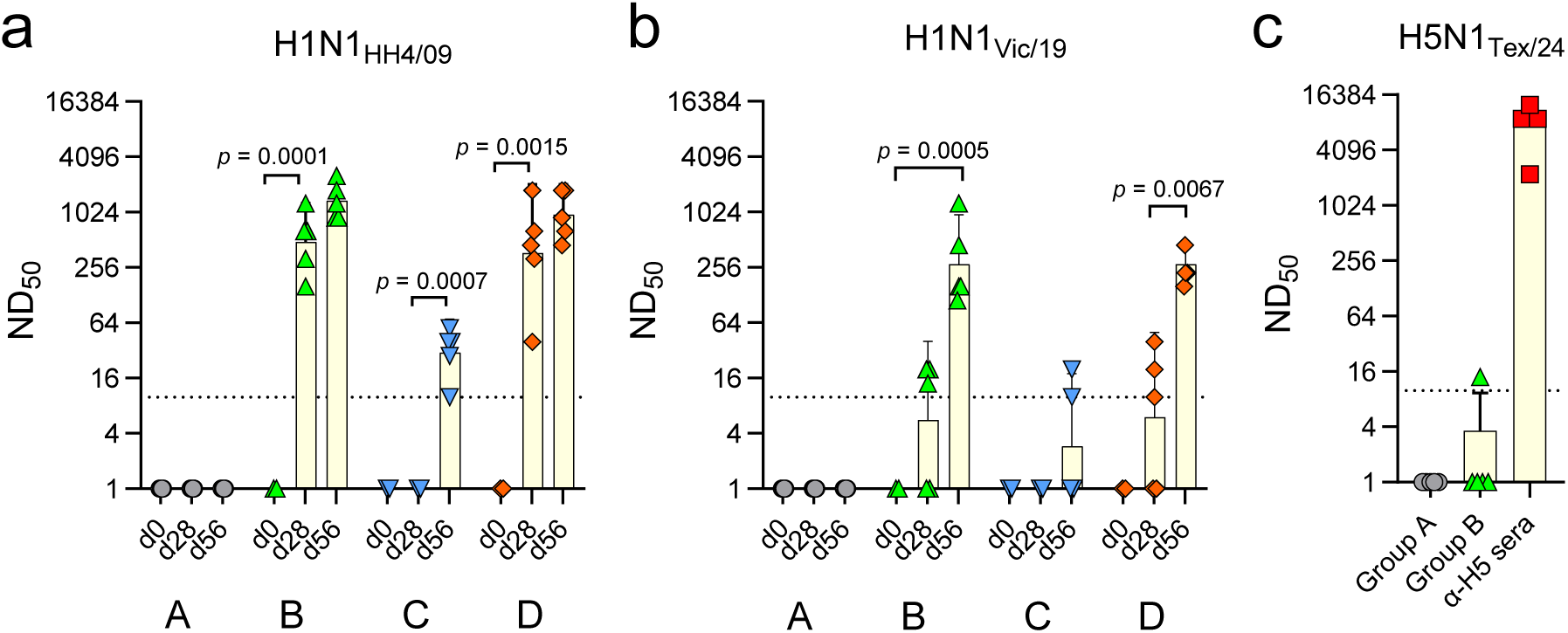
Determination of virus-neutralizing serum antibody levels in immunized pigs. SPF pigs were primed with either VSV*ΔG (groups A and C) or the VSVΔG(H1) + VSVΔG(N1) cocktail (groups B and D) and subsequently boosted with either VSV*ΔG (group A), VSVΔG(H1) + VSVΔG(N1) (group B) or the genetically modified LAIV (groups C and D). Serum was prepared prior to vaccination (d0), 4 weeks after the primary vaccination (d28), and 4 weeks after the second immunization (d56). **a, b** The virus-neutralization dose 50% (ND_50_) was determined against either H1N1_HH4/09_ (**a**) or H1N1_HH4/09_/H1N1_Vic/19_ 6:2 reassortant virus encoding the HA and NA antigens of H1N1_Vic/19_ (**b**). **c** The boost sera of groups A and B were tested for virus-neutralizing activity against H5N1_Tex/24_. Sera from chickens (n = 5) that had been vaccinated with VSVΔG(H5) encoding the HA of A/Dalmatian Pelican/Bern/1/2022 (H5N1) (α-H5) were used as positive control. Geometric mean values with 95% CI are shown. The two-way ANOVA test with Tukey’s multiple comparisons was used to identify significant ND_50_ values between pre-immune (d0), prime (d28), and boost (d56) sera.

### Vaccinated pigs produce N1-specific antibodies with sialidase-inhibitory activity

To measure the functional antibody response triggered by the VSVΔG(N1) vaccine component, we performed an enzyme-linked lectin assay (ELLA) to detect antibodies inhibiting the enzyme activity of the N1 neuraminidase. The assay is based on immobilized fetuin as substrate and biotin-labeled peanut agglutinin (PNA) as probe for the desialylated product of the enzyme reaction ^19,20^. Since the immune sera of immunized pigs also contain H1-specific antibodies that could sterically block the binding of N1-specific antibodies to the virion ^10^, the test was performed with H5N1_HH4/09_, a bovine H5N1 virus with its genome segment 6 (NA) replaced by the corresponding segment of H1N1_HH4/09_. For biosafety reasons, the proteolytic cleavage site in the H5 hemagglutinin was converted into a monobasic motif as found in low-pathogenic H5Nx influenza viruses ^21^. Pigs that had been vaccinated with the VSV*ΔG control vaccine (group A) did not develop antibodies with inhibitory activity against N1_HH4/09_ (**Fig. 3a**). Likewise, pigs immunized via the intranasal route with the LAIV (group C) did not develop significant levels of serum antibodies with N1 inhibitory activity. In contrast, pigs developed geometric mean sialidase inhibitory antibody titers of IC_50_ = 1267 (group B) and IC_50_ = 1283 (group D) after the prime immunization (i.m.) with the VSVΔG(H1) plus VSVΔG(N1) vaccine. The second intramuscular administration of the vaccine significantly boosted the N1-specific antibody response in group B (geom. mean IC_50_ = 7098), while the intranasal boost with the LAIV (group D) did not result in significantly increased inhibitory antibody titers (geom. mean IC_50_ = 2065).

**Fig. 3.**
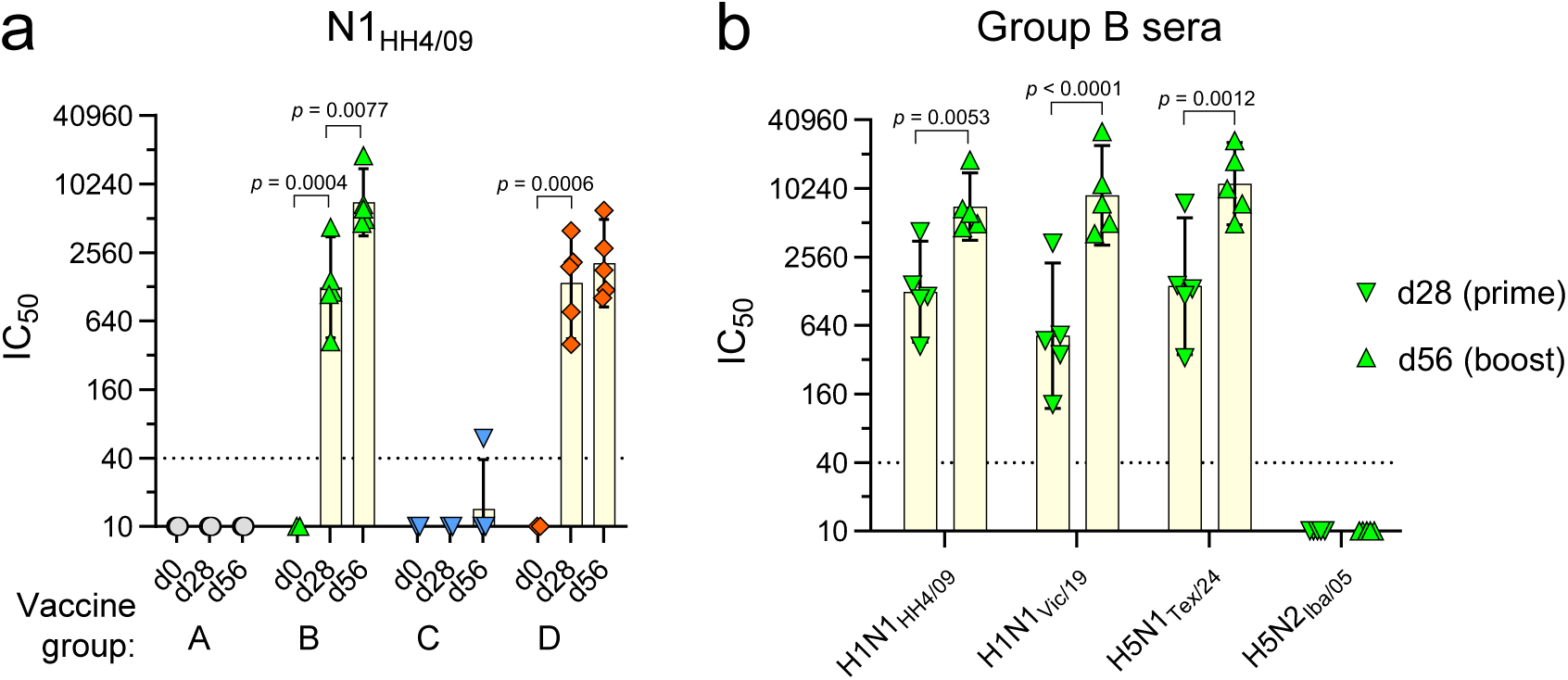
Detection of sialidase-inhibitory antibodies in sera of vaccinated pigs. The ELLA test was used to detect antibodies with inhibitory activity against the N1 sialidase of H5N1 reassortant viruses. **a** Preimmune (d0), prime (d28) and boost (d56) sera of the indicated vaccine groups were analyzed for inhibition of the N1_HH4/09_ sialidase. **b** Prime (d28) and boost (d56) sera of group B pigs were analyzed with H5N1 reassortant viruses encoding the NA of either H1N1_HH4/09_, H1N1_Vic/19_ or H5N1_Tex/24_, and H5N2_Iba/05_. The inhibitory concentration 50% (IC_50_) for each of the individual sera, geometric mean values (height of the bars) with 95% CI are shown. The two-way ANOVA test with either Sidak’s or Tukey’s multiple comparisons was used to assess significant differences between prime and boost sera and to compare the inhibition of different N1 neuraminidases. Only significant differences (p < 0.05) are indicated.

To analyze the inhibitory activity of the immune sera against N1 sialidases from other influenza viruses, we also generated H5N1 reassortant viruses encoding the N1 sialidases of either H5N1_Vic/19_ or H5N1_Tex/24_ and the N2 sialidase of A/chicken/Ibaraki/1/2005 (H5N2) (H5N2_Iba/05)_. Interestingly, the immune sera from group B pigs showed similarly high inhibitory activity against all three N1 sialidases but did not inhibit the N2 sialidase (**Fig. 3b**). These results show that the intramuscular immunization with the VSVΔG(N1) vaccine component led to the production of N1-specific serum antibodies that have potent sialidase inhibitory activity, even against distantly related N1 such as the one from the bovine H5N1_Tex/24_ virus.

### Antibodies to the N1_HH4/09_ neuraminidase inhibit the release of H5N1_Tex/24_ from infected MDCK cells

Immune sera from vaccinated pigs of group B did not show neutralizing activity against H5N1_Tex/24_ (see **Fig. 2c**) but showed potent inhibition of N1_Tex/24_ sialidase activity (see **Fig 3b**). To study the impact of N1 sialidase inhibition on the release of infectious H5N1_Tex/24_, we infected 96-well MDCK cell cultures with 100 TCID_50_/well of H5N1_Tex/24_ and then added cell culture medium containing serially diluted immune sera. These serum pools were prepared for all four vaccine groups four weeks after the second immunization. At 24 and 48 hours p.i., the infectious titers of H5N1_Tex/24_ released into the cell culture supernatants were determined on MDCK cells by end point titration. Immune sera of groups A and C did not inhibit the release of H5N1_Tex/24_ from infected MDCK cells, independently of the serum concentration used (**Fig. 4a, b**). In contrast, immune sera from group B showed a concentration-dependent inhibition of H5N1_Tex/24_ release. Using the immune serum at a dilution of 1:20, a remarkable reduction of infectious titer by 6 log_10_ was achieved at 24 h (**Fig. 4a**), which was still present at 48 h p.i. (**Fig. 4b**). Likewise, pooled group D immune serum mediated concentration-dependent inhibition of H5N1_Tex/24_ release, however, with general lower efficacy compared to group B immune serum. Analysis of the sera from individual animals allowed us to determine the serum concentration leading to 90% inhibition of H5N1_Tex/24_ release at 24 hours p.i. **(Fig. 4c).** The geometric mean IC_90_ titer was 896 for group B and 160 for group D, whereas group A and C sera had no significant impact on virus release. Release of a HPAI H5N4 virus was not inhibited by the group B serum pool, indicating that the inhibitory activity of the immune sera is NA subtype specific. At 48 h p.i., when the cell culture supernatant had been removed, the cells were fixed with formalin and infected cells visualized by indirect immunofluorescence using a monoclonal antibody directed to the influenza NP antigen. In MDCK cells incubated with group A or group C serum, H5N1_Tex/24_ infection spread throughout the whole monolayer, whereas in cells incubated with group B or D serum the infection remained restricted to distinct foci (**Fig. 4d**). Collectively, these results demonstrate that the VSVΔG(N1) vaccine component induces N1-specific serum antibodies with potent inhibitory activity against H5N1_Tex/24_.

**Fig. 4.**
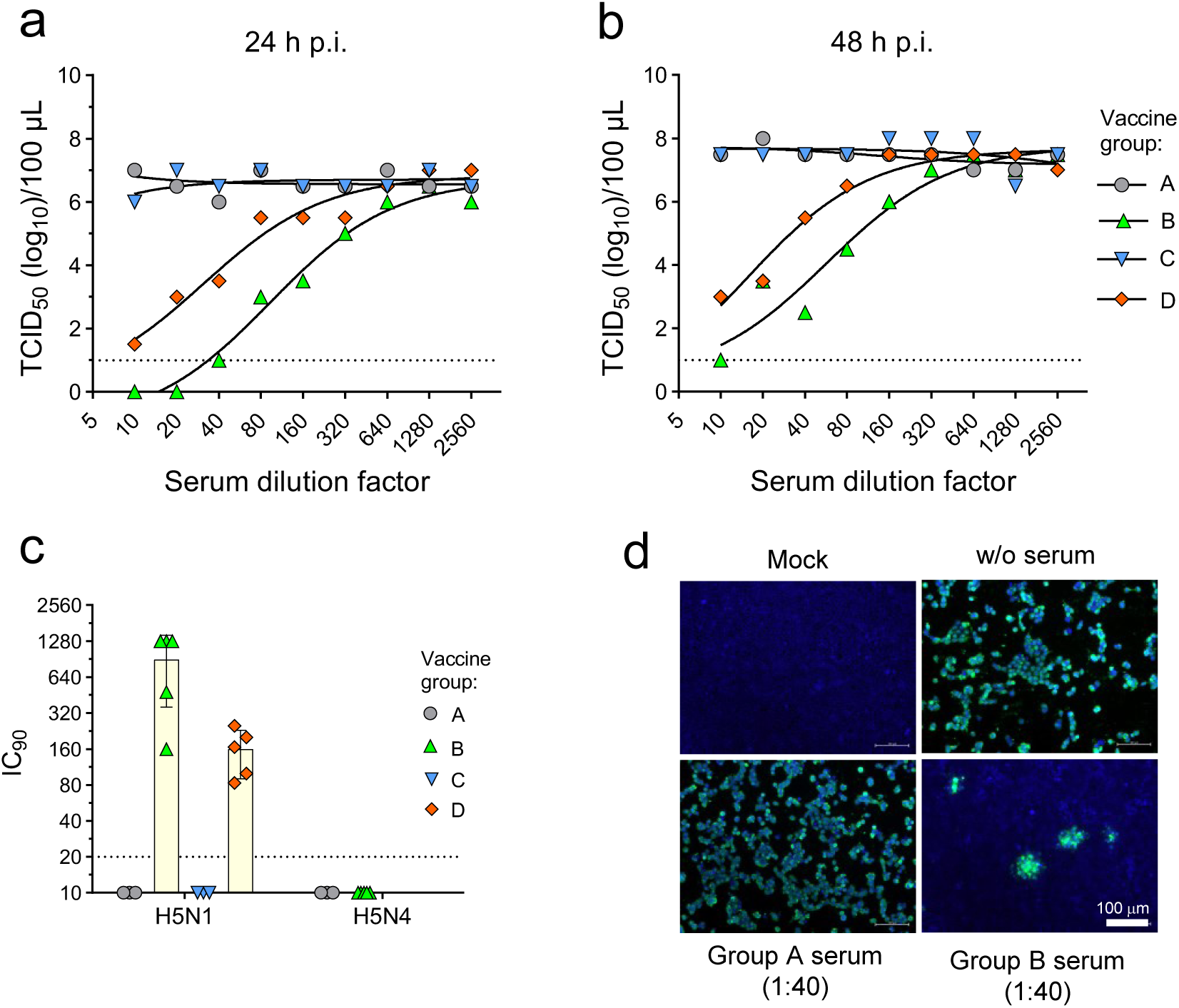
Antibodies raised against N1_HH4/09_ inhibit H5N1_Tex/24_ release from infected cells. MDCK cells grown in 96-well cell culture plates were inoculated for 60 min with 100 TCID_50_/well of H5N1_Tex/24_ and subsequently incubated with MEM containing serially diluted boost serum pools from the indicated vaccine groups. **a, b** Cell culture supernatant was collected at 24 h (**a**) and 48 h (**b**) post infection and infectious virus titers determined. The lower limit of detection is indicated by a dotted line. **c** Determination of the inhibitory concentration 90% (IC_90_) of boost sera from individual animals of the indicated vaccine groups causing 90% reduction of H5N1_Tex/24_ or H5N4_Swi/21_ release into the cell culture supernatant 24 h post infection. Only sera from group A and B animals were analyzed for inhibition of H5N4 release. The lower limit of detection is indicated by a dotted line. **d** At 48 h post infection with H5N1_Tex/24_, cells were fixed and the influenza NP antigen detected by indirect immunofluorescence (green fluorescence). Nuclei were stained by DAPI (blue fluorescence). Mock-infected cells and infected cells incubated with group A and group B sera at the indicated dilutions are shown. Bar size = 100 μm.

### Intramuscular priming of pigs eliminates LAIV shedding

In a recent study we showed that the prolonged shedding of the LAIV NS1(1-126)-ΔPAX from the upper respiratory tract of vaccinated pigs can be reduced if the animals were first primed with VSVΔG(H1) replicon particles ^22^. To confirm this finding and to see whether the vaccination of the pigs with VSVΔG(H1) in combination with VSVΔG(N1) could completely eliminate LAIV shedding, the vaccinated pigs of groups C and D were monitored for a total of 16 days by analyzing nasal swab samples by RT-qPCR for the presence of viral RNA (**Fig. 5a**) and infectious virus (**Fig. 5b**). Using these methods, we detected significant LAIV shedding by animals of group C, which lasted for several days. The three non-vaccinated pigs that were kept for three weeks with the animals of group C seroconverted to the influenza NP antigen (**Fig. 5c**), indicating that LAIV was transmitted to these sentinels. In contrast, if the pigs were first primed with the VSVΔG(H1)/VSVΔG(N1) vaccine, only some viral RNA was detected in nasal swabs at day 1 and 2 after nasal administration of the LAIV (**Fig. 5a**), and no shedding of infectious virus (**Fig. 5b**) or transmission to sentinels was detected (**Fig. 5c**). These data confirmed that intramuscular priming with the H1/N1 RNA replicon particles efficiently controlled replication of the LAIV in the upper respiratory tract.

**Fig. 5.**
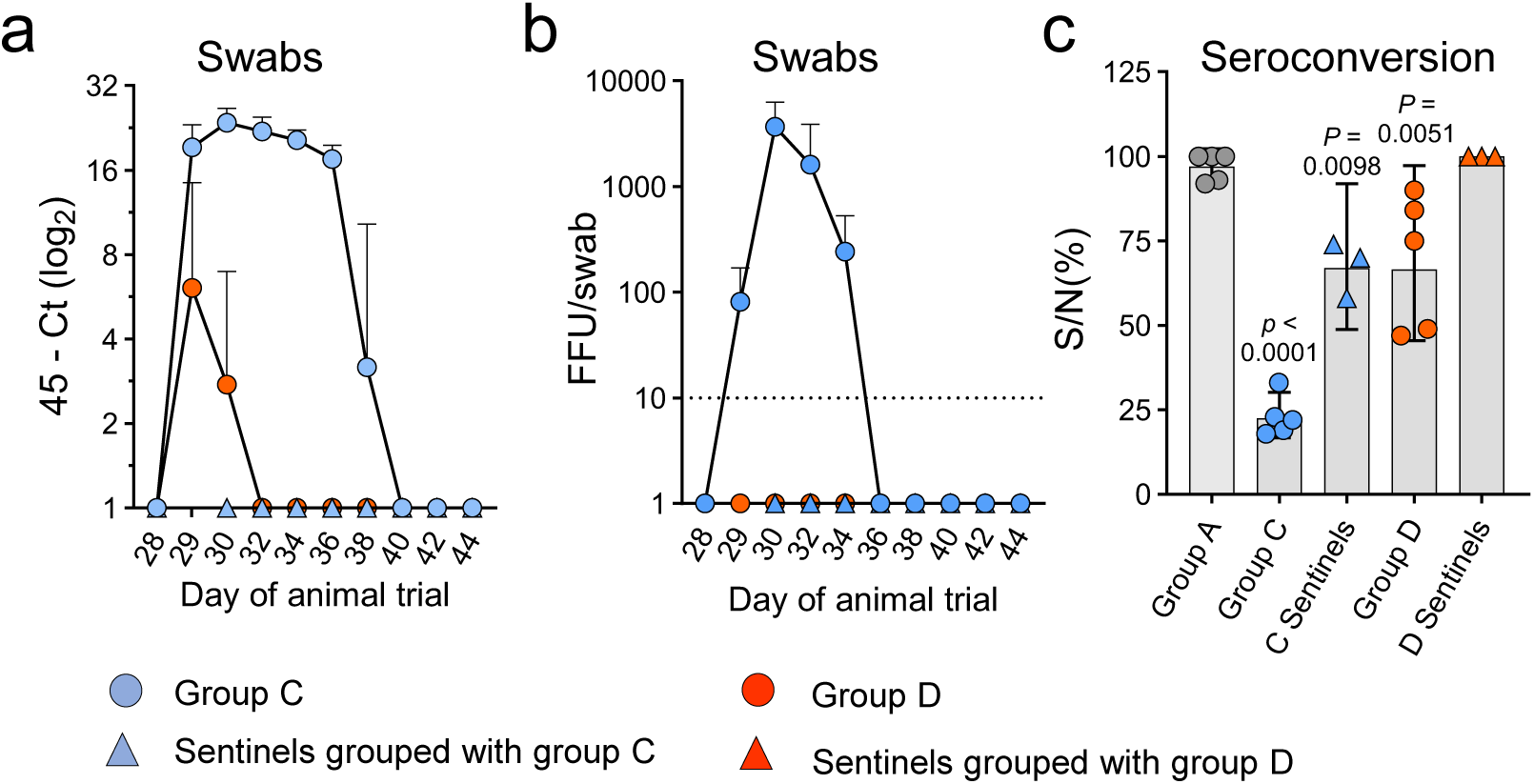
Analysis of LAIV shedding. Following primary immunization of group C and group D animals (n = 5 animals/group) with either VSV*ΔG (group C) or VSVΔG(H1)/VSVΔG(N1) cocktail (group D), the animals were boosted intranasally with the LAIV NS1(1-126)-ΔPAX. Sentinel animals (n = 3) were added to each group and co-housed with the LAIV boosted animals for three weeks. **a** RT-qPCR analysis of nasal swab samples collected at the indicated days post vaccination with LAIV. The 45-Ct is the cycle threshold number subtracted from the maximum number of cDNA amplification cycles. **b** Determination of the focus-forming units (FFU) in nasal swab samples suspended in 0.75 mL of MEM. The lower detection limit is indicated by a dotted line. **c** Analysis of seroconversion of pigs to the influenza NP antigen. Sera from group C and D animals were prepared four weeks after the booster vaccination (d56). Sera of the corresponding sentinel animals were prepared three weeks after they had been grouped with the vaccinated pigs (d49). NP-specific serum antibodies were detected by a competitive ELISA. The ratio of the optical density at 450 nm of sample to negative control (S/N%) was calculated. Group A animals that were immunized with the control vaccine VSV*ΔG served as negative control. A significant difference in S/N (%) indicating seroconversion was assessed by the one-way ANOVA test adjusted to Dunnett’s multiple comparisons and group A as reference.

### Vaccination of pigs efficiently prevents replication of antigen-drifted H1N1 virus in the respiratory tract

Four weeks after the second vaccination, the animals were challenged via the intranasal route with 10^6^ infectious units/pig of the H1N1_HH4/09_/H1N1_Vic/19_ 6:2 reassortant virus. None of the animals, including those in control group A, showed an elevated rectal temperature (**Supplementary Figure 1**) or other clinical symptoms (**Supplementary Tab. 2**) as a result of the infection. To monitor the shedding of challenge virus, nasal swab samples were collected daily for a total of 5 days post infection and analyzed for the presence of viral RNA (**Fig. 6a**) and infectious virus (**Fig. 6b**). As expected, all animals of control group A displayed shedding of challenge virus starting from day 2 and lasting until the end of the experiment at day 5 post infection (**Fig. 6a, b**). In contrast, no viral RNA was detected in swab samples taken from group C animals at any of the post challenge days, and only some swab samples collected from pigs of groups B and D were positive for viral RNA on some days (**Fig. 6a**). However, no infectious virus was detected in the nasal swabs collected from the pigs from groups B, C, and D (**Fig. 2b**). In line with these findings, viral RNA was detected in tonsils, mesenteric and tracheobronchial lymph nodes and lung lobes in most animals of group A but was essentially absent from the corresponding tissues of animals of group B, C, and D **(Fig. 6c).** These results show that both the heterologous and homologous prime/boost vaccination regimens are effective in preventing the replication and shedding of antigen-drifted virus.

**Fig. 6.**
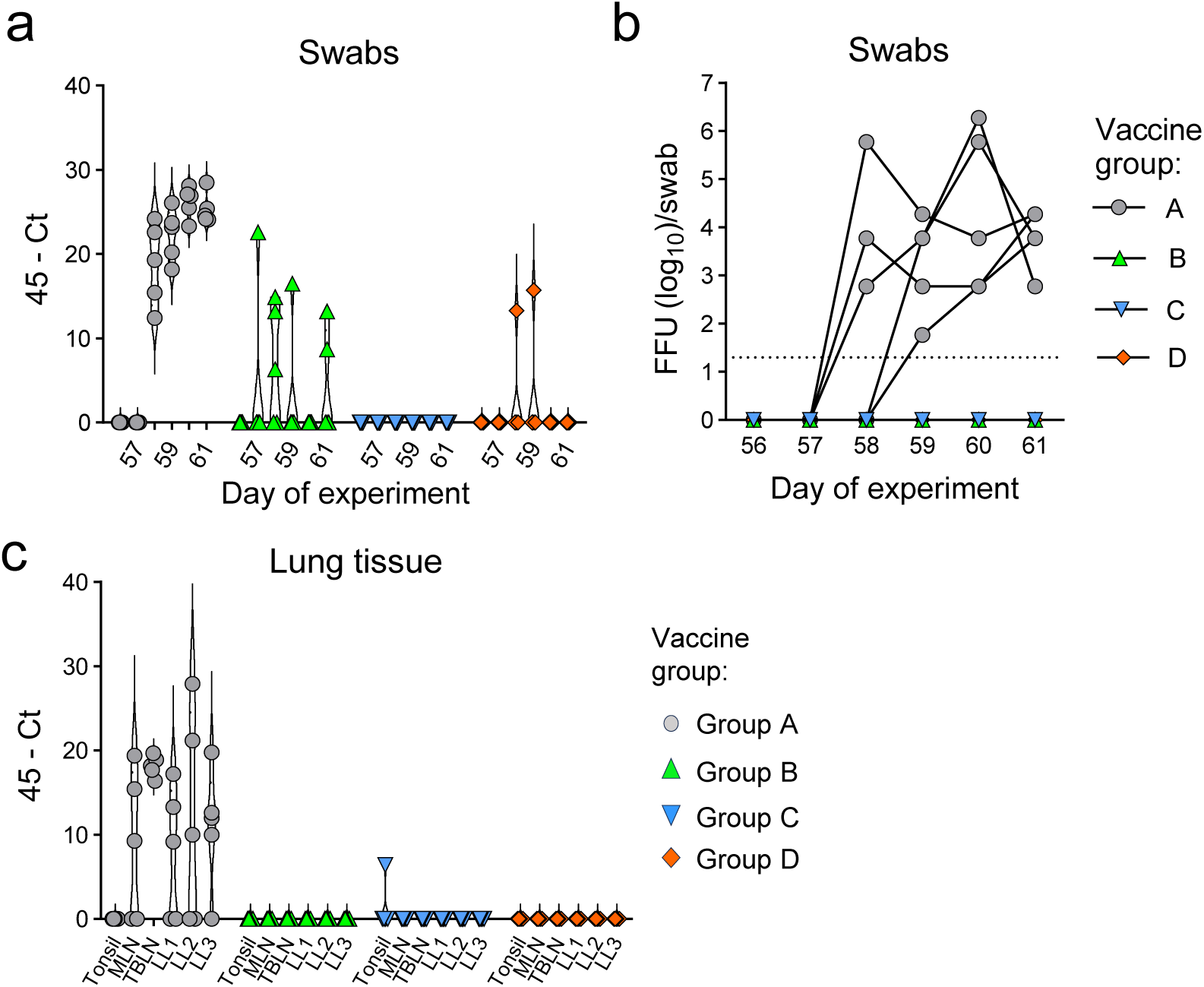
Virus loads in lung tissue and nasal swab samples. Four weeks after the booster vaccination, SPF pigs were challenged via the nasal route with the H1N1_HH4/09_:H1N1_Vic/19_ 6:2 reassortant virus encoding the HA and NA antigens of H1N1_Vic/19_. **a** Detection by RT-qPCR of viral RNA in nasal swab samples collected at the indicated days post infection. The cycle threshold (Ct) subtracted from 45, i.e. the maximum number of cycles performed, is shown. **B** Detection of infectious virus in nasal swab samples. Swab samples were suspended in 0.75 mL of MEM and infectious virus titrated on MDCK cells. The focus-forming units (FFU) are shown for all swabs collected from individual animals in the indicated groups. The lower detection limit is indicated by a dotted line. **c** Detection by RT-qPCR of viral RNA extracted from tonsils, mesenteric lymph nodes (MLN) tracheobronchial lymph nodes (TBLN), lung lobes (LL) of euthanized pigs at day 5 post infection.

## Discussion

### Heterologous prime/boost immunization

Antigen-drift represents a major hurdle for the efficacy of conventional influenza vaccines that rely on inactivated split viruses and are standardized to the amount of purified HA antigen. In the present work, we show that intranasal immunization of pigs (group C) with the LAIV NS1(1-126)-ΔPAX triggers an immune response that efficiently controlled the replication of a reassortant virus bearing the HA and NA envelope glycoproteins of the antigen-drifted H1N1_Vic/19_ strain. Although intranasal immunization with LAIV induced relatively low levels of virus-neutralizing serum antibodies, previous work with the same LAIV showed that intranasal immunization of pigs with this vaccine induces local immune responses including secretion of H1N1-specific IgA into the mucosal tissues and cellular immune responses that are expected to contribute to its potent inhibition of challenge virus replication in the mucosa ^22^. However, the prolonged shedding of LAIV via the respiratory tract and transmission to sentinel pigs was identified as a safety issue ^22–24^. This problem was solved by first immunizing the animals via the intramuscular route with VSV-based RNA replicon particles encoding HA ^22^. This immune priming induced high levels of systemic immunity that efficiently controlled LAIV shedding without preventing the LAIV from boosting neutralizing antibodies. In the present study, immune priming with both H1 and N1 antigen also eliminated LAIV shedding. Still, the boost vaccination using the LAIV significantly enhanced the systemic immune response against the antigen-drifted HA_Vic/19_ protein. Furthermore, the inclusion of VSV replicons encoding the N1_HH4/09_ antigen resulted in the induction antibodies that inhibited the sialidase activity of N1_HH4/09_ and N1_Vic/19_ equally well. As a result, this vaccination protocol induced robust cross-protective immunity, even preventing shedding of heterologous challenge virus from the respiratory tract.

### Homologous prime/boost immunization

The first immunization (i.m.) with the combined VSVΔG(H1) and VSVΔG(N1) vaccine resulted in high antibody titers with neutralizing activity against homologous H1N1_HH4/09,_ which increased only moderately after administration of the second vaccine dose. In contrast, the neutralizing activity against antigenically drifted H1N1_Vic/19_ strain was strongly boosted by the second dose. This broadened immunity was likely driven by memory B cells undergoing affinity maturation upon antigen re-exposure ^25,26^. Notably, re-stimulating memory B cells with the same antigens enhances not only the strength and quality but also the longevity of antibody responses ^27^.

Another important finding of our study was that homologous prime/boost strategy using the combined replicon particle vaccines eliminated the shedding of the antigen-drifted H1N1 challenge virus, even though the vaccines were administered via the intramuscular route. ^28^. Possibly, H1- and N1-specific serum IgG was secreted into the mucosal tissues of the respiratory tract via the neonatal Fc receptor (FcRn) which is expressed in mammalian species also at adult life ^29^. In mice, passive intravenous transfer of anti-influenza immunoglobulins indicated that high serum IgG levels can efficiently reduce nasal influenza virus shedding ^28^, suggesting that a fraction of the serum IgG can be secreted to mucosal surfaces. Thus, the strong immune response induced by intramuscular immunization with the RNA replicon particles is the likely key factor preventing challenge virus replicating in the respiratory mucosa and, consequently, virus shedding.

### Importance of NA-specific antibodies in protection

In contrast to our previous study in which the VSVΔG(H1) vaccine was used for immunization of pigs ^22^, the present work made also use of VSVΔG(N1) replicon particles encoding N1_HH4/09_. The N1-specific antibodies elicited by this vaccine potently inhibited the sialidase activity of the homologous N1_HH4/09_ but also the sialidase activity of N1_Vic/19_ and N1_Tex/24_, although the latter shows only 88% amino acid identity with N1_HH4/09_ (**Supplementary Fig. 2**). This suggests that the N1-specific antibodies bound to epitopes that are conserved in this antigen. There is evidence that the antigenic drift associated with NA is slower than that of the HA antigen, possibly because of functional constraints, i.e. only mutations that do not significantly impair sialidase activity are tolerated ^12,30^.

NA sialidase activity is crucial for the efficient release of progeny influenza virus from the infected host cell ^31,32^ but also helps the virus to penetrate the mucus layers covering the epithelial cells of the respiratory tract ^33,34^. Thus, the antibodies induced by VSVΔG(N1) likely contributed to immune protection of the pigs by inhibiting virus release and spread. Indeed, previous work has shown that VSVΔG(NA) replicon particles are protective even if used as a stand-alone vaccine ^10,35^. The observation that immune sera of pigs immunized with the combined VSVΔG(H1) and VSVΔG(N1) replicon particles potently inhibited release and spread of H5N1_Tex/24_, despite lacking virus-neutralizing activity, underscores the enhanced broad cross-reactivity achieved by targeting NA in vaccination.

In recent years, H5N1 HPAI viruses of the clade 2.3.4.4b have spread worldwide and led to the death of millions of birds of numerous species ^36^. In addition, clade 2.3.4.4b viruses showed the remarkable potential to infect various mammalian species ^37,38^, including humans ^39^. While infection of cats by H5N1 caused neurological symptoms with often fatal outcome ^40^, infection of humans was mostly associated with mild symptoms such as conjunctivitis ^41^. There is evidence that infection with human influenza viruses elicits broadly cross-reactive NA-specific antibodies that protect from infection by HxN1 viruses *in vitro* and *in vivo* ^42,43^. The protective effect of prior infection with 2009 pandemic H1N1 virus on the pathogenesis and transmission of clade 2.3.4.4b H5N1 virus was also observed in the ferret model ^44,45^. These findings imply that children who have not experienced infection with seasonal H1N1 virus yet might be at higher risk to develop severe symptoms if infected with H5N1 HPAI. Indeed, severe disease following infection with clade 2.3.4.4b genotype D1.1 was documented for a 13-year-old teenager in 2024 in Canada ^46^ and a fatal infection of a 3-year-old girl was reported in 2025 in Mexico ^47^. However, it is not known whether these children were more susceptible to H5N1 infection because they had no pre-existing immunity to influenza viruses in general.

Taken together, the present study not only confirms the efficacy and safety of influenza vaccines based on non-replicating RNA replicon particles in the porcine animal model ^22,48,49^, but demonstrates that the high levels of neutralizing anti-HA as well as the NA-targeting antibodies ensure broad systemic and mucosal protection against antigen-drifted influenza viruses. These properties represent a significant improvement compared to currently used inactivated seasonal influenza vaccines ^50,51^.

## Methods

### Cells

Madin-Darby canine kidney type II cells (MDCK-II) were kindly provided by Georg Herrler (University of Veterinary Medicine, Hannover, Germany) and maintained with minimum essential medium (MEM, Thermo Fisher Scientific, Basel, Switzerland; cat. no. 31095-029) supplemented with 5% of fetal bovine serum (FBS; Pan Biotech, Aidenbach, Germany; cat. no. P30-3033). Human embryonic kidney (HEK) 293T cells (American Type Culture Collection (ATCC), Manassas, USA; cat. no. CRL-3216) were maintained with Dulbecco’s Modified Eagle Medium (DMEM, Thermo Fisher Scientific; cat. no. 32430-027) supplemented with 10% FBS. Baby hamster kidney 21 (BHK-21) fibroblasts were obtained from ATCC (cat. no. CCL-10) and maintained in Glasgow’s minimal essential medium (GMEM, Thermo Fisher Scientific, cat. no. 21710-025) supplemented with 5% FBS. BHK-G43 cells, a transgenic BHK-21 cell clone expressing VSV glycoprotein G in a regulated manner ^52^, was maintained in GMEM supplemented with 5% FBS. All cells were grown at 37 °C in a 5% CO_2_ atmosphere.

### Viruses

The HPAI virus A/cattle/Texas/063224-24-1/2024 (H5N1) (H5N1_Tex/24_), clade 2.3.4.4b, genotype B3.13 (GISAID accession number: EPI_ISL_19155861), was kindly provided by Diego Diel (Cornell University, Ithaca, NY, USA). This virus was originally isolated from the milk of infected dairy cows in Texas, USA ^53^. A/Buzzard/Switzerland/V0052/2021 (H5N4) (H5N4_Swi/21_) was isolated by IVI Mittelhäusern from a brain biopsy sample taken from a common buzzard that was found dead close to Lake Constance in 2021. The 2009 pandemic virus strain A/Hamburg/4/2009 (H1N1) (H1N1_HH4/09_) was produced by reverse genetics using the 8-plasmid system ^15^. The live-attenuated influenza vaccine (LAIV) NS1(1-126)-ΔPAX is based on a modified H1N1_HH4/09_ and encodes a truncated NS1 protein and is unable to drive expression of the accessory PA-X protein ^15^. VSV*ΔG is a propagation-defective vesicular stomatitis virus in which the glycoprotein (G) gene has been replaced by the green fluorescent protein (GFP) gene ^54^.

### Generation of recombinant influenza viruses

The cDNAs encoding the genomic segments 4 and 6 of A/Victoria/2570/2019 (H1N1) (GenBank acc. nos. WEY08940 and WEY08939, respectively) and the cDNAs of the eight genomic segments of A/bovine/Texas/24-029328-02/2024 (H5N1) (GenBank accession nos. PP599470–PP599477) ^55^ were synthesized by Genscript Biotech (Piscataway, New Jersey, USA). The synthetic cDNA of RNA segment 4 encoded an HA containing the monobasic cleavage site P_337_LRETR↓GLF rather than the polybasic cleavage site P_337_LREKRRKR↓GLF found in the authentic virus. All cDNAs were amplified with Phusion DNA and cloned into the *Bsm*BI restriction site of the pHW2000 plasmid ^54^ using the In-Fusion cloning system (Takara Bio, Saint-Germain-en-Laye, France, cat. no. 638948). The primary nucleotide sequence of each of the cloned segments was verified by Sanger sequencing.

Infectious viruses were generated by transfection of HEK 293T/MDCK cell co-cultures using the respective eight genomic cDNAs cloned into the pHW2000 vector and Lipofectamine 2000 (Life Technologies, cat. no. 11668019) as transfection reagent. In this way, the following recombinant viruses were generated: A virus containing segments 4 and 6 of H1N1_Vic/19_ and the remaining six segments of H1N1_HH4/09_, a H5N1_Tex/24_ with a monobasic proteolytic cleavage site in HA, and three reassortant viruses encoding segment 6 of either H1N1_HH4/09_, H1N1_Vic/19_, or H5N1_Tex/24_ and the remaining genomic RNA segments of (H5_mb_N1)_Tex/24_. The rescued viruses were passaged two times on MDCK cells with FBS-deficient medium containing 1% penicillin/streptomycin and 1 µg/mL of acetylated trypsin (Merck KGaA, Darmstadt, Germany, cat. no. 4370285). Virus stocks were stored in aliquots at −70 °C in the presence of 5% FBS.

### Titration of influenza A viruses

For titration of IAV that had a monobasic cleavage site in their HA, MDCK cells grown in 96-well tissue culture plates were inoculated in duplicate with 40 µL per well of serial 10-fold virus dilutions for 1 h at 37 °C. Thereafter, 160 µL of MEM containing 1% methylcellulose were added to each well, and the cells incubated for 24 h at 37 °C. The cells were fixed for 30 min with 4% formalin in PBS, permeabilized with 0.25% (v/v) of Triton X-100, and incubated for 60 min with a monoclonal antibody directed to the influenza nucleoprotein (NP) antigen (American Type Culture Collection, Manassas, Virginia, USA, ATCC HB-65, clone H16-L10-4R5, diluted 1:50 in PBS), and subsequently for 60 min with goat anti-mouse IgG conjugated with Alexa Fluor-488 (Life Technologies, Zug, Switzerland; diluted 1:500 in PBS). Infected cells were detected by fluorescence microscopy (Observer Z1 microscope, Zeiss, Feldbach, Switzerland), and infectious virus titers were calculated and expressed as focus-forming units per milliliter (ffu/mL).

For titration of the HPAI H5N1_Tex/24_, MDCK cell monolayers in 96-well tissue culture plates were incubated in quadruplicates with serially diluted virus (100 μL/well) for 48 h at 37 °C. The cells were washed once with PBS (200 μL/well) and fixed for 1 h at 21 °C with 4% of buffered formalin containing 0.1% (w/v) crystal violet (Merck KGaA, cat. no. 1.01408). The plates were washed with tap water and dried. Virus titer was calculated using the Spearman-Kärber method and expressed as tissue culture infectious dose 50% (TCID_50_) ^56^.

### Generation of recombinant VSV vector vaccine

For generation of the VSVΔG(H1) and VSVΔG(N1) vaccine components, the HA and NA genes of H1N1_HH4/09_ (GenBank acc. nos. GQ166213 and GQ166217, respectively) were amplified by PCR using the Phusion DNA polymerase and inserted into the *Mlu*I and *Nhe*I sites of the pVSV*ΔG(HA_H5-HP_) plasmid ^57^, resulting in the pVSVΔG(H1) and pVSVΔG(N1) plasmids, respectively. VSVΔG(H1) and VSVΔG(N1) replicon particles were generated from transfected cDNA ^49,58,59^ and propagated on BHK-G43 cells expressing the VSV G protein in a regulated manner ^52^. Virus stocks were stored in aliquots at −70 °C. Infectious virus titers were determined on BHK-21 cells by end point dilution. Infected cells were detected 20 h p.i. by indirect immunofluorescence using a monoclonal antibody (mAb 23H12) directed to the VSV matrix protein (KeraFast, Boston, MA, cat. no. EB0011) ^60^.

For production of vaccine batches, BHK-G43 cells were seeded into five T150 flasks and maintained in 40 mL/flask of GMEM medium with 5% FBS. When confluency was reached, the medium was aspirated and the cells maintained at 37 °C for 6 h with 40 mL/flask of GMEM containing 10^−9^ M of mifepristone. VSVΔG(H5_mb_) was added at a multiplicity of infection (moi) of 0.05 ffu/cell and incubated with the cells for 20 h at 37 °C. The cell culture supernatant was transferred to 50-mL Falcon tubes and cell debris removed by centrifugation (1000 × g, 15 min, 4 °C). The replicon particles were pelleted from the cleared supernatant by ultracentrifugation (105’000 × g, 60 min, 4 °C) and resuspended in 20 mL of PBS. The VSVΔG(H1) and VSVΔG(N1) replicon particles were stored in aliquots at −70 °C.

### Animal experiments

The animal experiments were performed according to Swiss laws (the Animal Welfare Act TSchG SR 455, the Animal Welfare Ordinance TSchV SR 455.1, and the Animal Experimentation Ordinance TVV SR 455.163). All experiments were reviewed by the committee on animal experiments of the canton of Bern and were approved by the cantonal veterinary authority under license number **BE15/2024**.

Twenty 10-week-old specific pathogen-free (SPF) Swiss Large White pigs from the breeding facility of the Institute of Virology and Immunology IVI (Mittelhäusern, Switzerland) were randomly divided into four groups (A – D), each containing five pigs of both sexes (n = 5). After an acclimatization period of five days, the animals of groups A and C received 2 mL of VSV*ΔG suspension (4 × 10^8^ ffu/mL) that were injected into the *m. brachiocephalicus* (neck muscle) (**Fig. 1**). The animals of groups B and D were immunized by injection of 2 mL (4 × 10^8^ ffu/mL) of VSVΔG(H1) and 2 mL (4 × 10^8^ ffu/mL) of VSVΔG(N1) suspension into different sites of the neck muscle. Body temperature and clinical symptoms were monitored daily for seven days, and serum samples were prepared at day 28 and day 56 after primary immunization (**Fig. 1**). At day 28, the animals of group A received a second dose (2 mL, 4 × 10^8^ ffu/mL) of the control vector VSV*ΔG while animals of group B were immunized (i.m.) with VSVΔG(H1) and VSVΔG(N1) replicon particles using the same doses as for the first time. Animals of groups C and D were immunized via the nasal route with 5 mL of PBS containing 2 × 10^4^ ffu/mL of LAIV. To this end, an intranasal mucosal atomization device (MAD Nasal, Teleflex Medical Europe Ltd., Ireland, cat. no, MAD300) plugged to a 10-mL syringe was employed.

After the first and the second immunization, the rectal temperature of the pigs and potential vaccine-mediated adverse effects were recorded daily for a total of 10 days. Blood (10 mL) was collected prior to vaccination (day 0), 28 days after prime vaccination, and 28 days after the booster immunization (day 56). Serum was prepared after coagulation of the blood and stored at −20 °C prior to analysis by virus neutralization, sialidase inhibition and virus release inhibition tests. To monitor shedding of LAIV, two nasal swabs were taken from each animal of groups C and D prior to vaccination (day 28) and at days 29, 30, 32, 34, 36, 38, 40, 42, 44. One swab sample was soaked in 0.75 mL of MEM supplemented with penicillin/streptomycin (Life Technologies, cat. no. 15140-122) and amphotericin B (Life Technologies, cat. no. 15290018), while the second swab was soaked in 0.75 mL of RA1 lysis buffer (Macherey-Nagel, Düren, Germany; cat. no. 740961) containing 1% (v/v) β-mercaptoethanol. Swab-soaked media and lysates were stored at −70 °C prior to analysis by virus titration and RT-qPCR, respectively. To monitor transmission of the LAIV, three sentinel animals were co-housed with each group C and group D at day 30 of the experiment (48 h after the second immunization). The sentinel animals were euthanized at day 54 and their serum tested for the presence of NP-specific antibodies by ELISA.

On day 56, animals in groups A to D were infected intranasally with 2 ml PBS containing 10^6^ ffu/ml of the H1N1_HH4/09_ /H1N1_Vic/19_ 6:2 reassortant. After infection, rectal temperature and clinical symptoms, including liveliness, body tension, body shape, appetite, nasal discharge, respiratory rate, expiratory effort, wheezing/coughing and diarrhea, were recorded daily for a total of 5 days. A clinical scoring system was used to assess disease severity and determine the humane endpoint: 0 = no symptoms, 1 = mild symptoms, 2 = moderate symptoms, 3 = severe symptoms. The animals were euthanized immediately if they reached a total clinical score of 9, or if a clinical score of 3 was reached for one or more of the clinical parameters. In addition, nasal swab samples were taken daily and tested for the presence of viral RNA using RT-qPCR. Five days after infection, the animals were killed by electrocution and subsequent exsanguination. Immediately after exsanguination, biopsy material was taken from the tonsils, mesenteric lymph nodes, tracheobronchial lymph nodes and the right cranial lobe of the lung and tested for the presence of viral RNA using RT-qPCR.

### Sialidase inhibition assay

The heat-inactivated immune sera of all animals in a group were mixed to obtain group-specific serum pools for both the primary and the booster vaccinations. Starting with a dilution of 1:20, four-fold serially diluted serum pools were prepared in 96-well plates using PBS as diluent. To 50 μL of each serum dilution, 50 μL of NA reassortant H5N1 virus (10^7^ ffu/mL) were added (in duplicates) and incubated for 30 min at 37 °C, The serum-virus mixture (100 μL) was then transferred to Nunc MaxiSorp 96-well plates (Thermo Fisher Scientific) that had been coated for 24 h at 4 °C with 10 µg/well of bovine fetuin (Merck KGaA, Darmstadt, Germany, cat. no. F2379) and subsequently blocked with 1% bovine serum albumin (BSA, Fluka Chemie GmbH, Buchs, Switzerland, cat. no. 05480). The microtiter plates with the immobilized fetuin were incubated with the serum-virus mixtures for 20 h at 37 °C and then washed three times with 250 μL/well of PBS containing 0.05% Tween 20 (PBS-T). Subsequently, 50 μL/well of PBS containing 0.1% BSA and 0.5 µg/mL of biotinylated peanut agglutinin (PNA, Merck KGaA cat. no. L6135) were added and incubated for 1 h at 21 °C. The microtiter plates were washed three times with PBS-T before 50 µL/well of streptavidin-peroxidase conjugate (Dako, Glostrup, Denmark; 1:5000 in PBS) were added and incubated with the plates for 30 min at 21 °C. The wells were washed as above again and 50 µL of 3,3′,5,5′-Tetramethylbenzidine (TMB) peroxidase substrate (Merck KGaA, cat. no. T4444) was added to each well. The reaction was stopped by adding 50 μL/well of 1 M hydrochloric acid (Merck KGaA, cat. no. 1.09057.1000) and absorption was measured at 450 nm using the GloMax® Discover microplate reader (Promega Corporation, Madison, WI, USA). The OD_450_ values were plotted against the serum dilution factors, and the dilution factor resulting in 50% inhibition (inhibitory concentration 50% [IC_50_]) was calculated by non-linear regression analysis (curve fitting).

### Virus neutralization test

Porcine sera were heated for 30 min at 56 °C to inactivate complement factors. Serial two-fold dilutions in MEM medium were added in quadruplicates to 96-well microtiter plates (50 µL/well). To each well, 50 µL of H1N1_HH4/09_, H1N1_Vic/19_ or H5N1_Tex/24_ (2000 ffu/mL) were added and incubated for 1 h at 37 °C. The antibody/virus mix was incubated for 1 h at 37 °C and 5% CO_2_ with MDCK-II cells that were grown to confluence in 96-well cell culture plates. Thereafter, the cells were washed once and incubated for 24 h at 37 °C with 100 µL/well of MEM medium. The cells were fixed with 4% formalin and permeabilized with PBS containing 0.25% Triton X-100. Virus-infected cells were visualized by indirect immunofluorescence using a monoclonal antibody directed to the influenza nucleoprotein (NP) antigen (American Type Culture Collection, Manassas, Virginia, USA, ATCC HB-65, clone H16-L10-4R5, diluted 1:50 with PBS) and goat anti-mouse IgG conjugated with Alexa Fluor-488 (Life Technologies, Zug, Switzerland; diluted 1:500 in PBS) ^61^. The 50% neutralizing dose (ND_50_) was calculated using the Spearman-Kärber method ^56^.

### Virus release inhibition assay

MDCK cells were grown in 96-well tissue culture plates and inoculated for 30 min at 37 °C with 50 µl MEM containing 100 ffu of H5N1_Tex/24_. The cells were washed once with 200 µL of Dulbeccòs phosphate-buffered saline (DPBS; Life Technologies, cat. no. 14040-091) and incubated at 37 °C with 200 uL of MEM cell culture medium containing serially two-fold diluted immune sera (boost) and supplemented with penicillin and streptomycin (Life Technologies, cat. no. 15140-122). At 24 h and 48 h post infection, 100 µL aliquots of cell culture supernatant were collected and frozen at −70 °C. The number of infectious virus particles released was determined as described above (see section “Titration of influenza A viruses”).

### RT-qPCR

Swab samples were directly transferred to 700 µl of RA1 lysis buffer (Macherey-Nagel, Düren, Germany, cat. no. 740961) containing 1% β-mercaptoethanol. Organs were homogenized in RA1 lysis buffer using a tissue bullet blender (Next Advanced Inc., Troy, NY, USA). Total RNA was extracted from the lysates using the NucleoMag Vet kit (Macherey-Nagel, cat.no. 744200) according to the manufacturer’s protocol. Reverse transcription from RNA to cDNA and real-time quantitative PCR (qPCR) were performed on a QuantStudio 5 real-time PCR system (Thermo Fisher Scientific) using the AgPath-ID One-Step RT-PCR kit (Life Technologies, Zug, Switzerland, cat. no. AM1005) with vRNA segment 7-specific oligonucleotide primers and probe ^62,63^. Data were acquired and analyzed using the Design and Analysis Software v1.5.2 (Thermo Fisher Scientific).

### Statistical analysis

Statistical analyses were performed using GraphPad Prism 10, version 10.1.2 (GraphPad Software, Boston, Massachusetts, USA). Unless noted otherwise, data are presented as scatter dot plots with geometric mean values and 95% confidence interval (CI) also indicated. Specific statistical tests such as the one-way or two-way ANOVA test were used to assess significant differences in serum antibody responses in vaccinated animals as indicated in the figure legends. *P* values < 0.05 were considered significant.

## Acknowledgements

We like to thank the Swiss National Research Foundation (SNSF) and the European Union’s Horizon Europe Project 101136346 EUPAHW for financial support of this project. We are grateful to Daniel Brechbühl and Katarzyna Sliz for their support in animal experimentation. We like to thank Martin Schwemmle (University of Freiburg, Germany) and Yoshihiro Sakoda (University of Sapporo, Japan) for providing plasmids and Diego Diel (Cornell University, USA) for providing the bovine H5N1 virus isolate.

## Author contributions

AS and GZ conceived the study and were responsible for funding acquisition; NR, OG and GZ conceptualized and conducted the animal experiments; RA, LB, and GZ generated recombinant influenza viruses and VSV replicon vaccines; LB performed RT-qPCR analysis, virus neutralization tests, virus release inhibition tests, and sialidase inhibition tests; LB and GZ analyzed and evaluated the data. GZ wrote the manuscript draft. All authors read and approved the final manuscript.

## Data availability

All data generated or analyzed during this study are included in this published article and its supplementary information files. Raw data are available at Zenodo (https://doi.org/10.5281/zenodo.17302034).

## Competing interests

RA, GZ, and AS filed a patent application related to the intramuscular prime/intranasal boost vaccine described in this work. All other authors declare no competing interests.

## Funding

This work has received funding from the Swiss National Research Foundation (SNSF), grant no. 189903 (A.S., G.Z.; https://data.snf.ch/grants/grant/189903) and was co-funded by the European Union’s Horizon Europe Project 101136346 EUPAHW by a grant to A.S.. The funders had no role in study design, data collection and analysis, decisions to publish or preparation of the manuscript.

---

**Supplementary Fig. 1.**
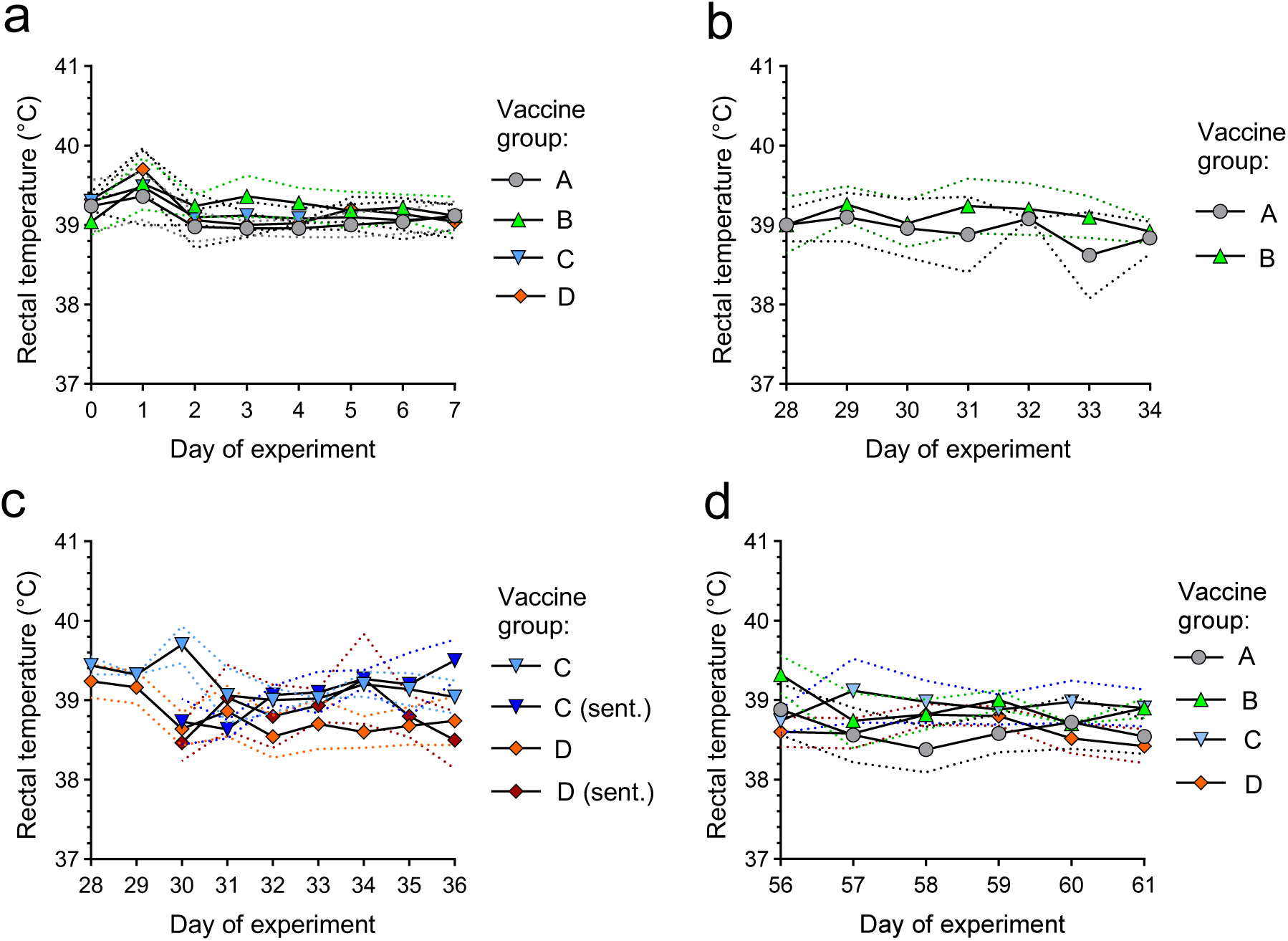
Recording of the rectal temperature in SPF pigs following vaccination and challenge infection. **a-d** The rectal temperature was recorded in all animals of the indicated groups (n = 5 animals/group) at the indicated day of the experiment. Mean values with SD (dotted lines) are shown. **a** Recording of rectal temperature following prime vaccination of groups A - D. **b** Recording of the rectal temperature in animals of groups A and B following boost vaccination. **c** Recording of rectal temperature in animals of group C and D (n = 5/group) after receiving the LAIV, and in the sentinel pigs (n = 3) that were co-housed with each of the groups. **d** Recording rectal temperature in pigs of all groups following nasal challenge infection with the H1N1_HH4/09_/H1N1_Vic/19_ 6:2 reassortant virus.

**Supplementary Fig. 2.**
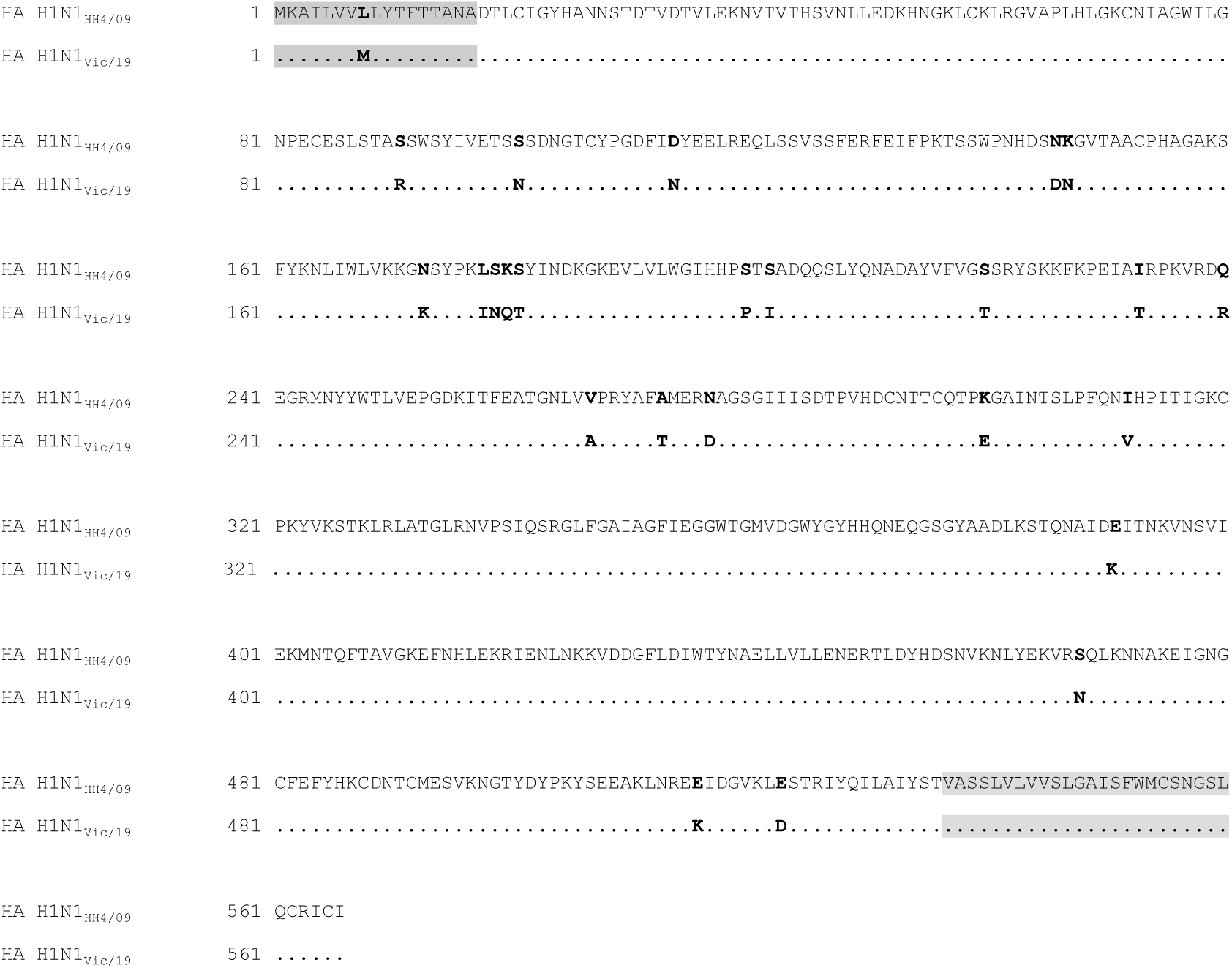
Alignment of HA primary amino acid sequences. The HA sequences of A/Hamburg/4/2009 (H1N1) (GenBank accession no. ACR10223) and A/Victoria/2570/2019 (H1N1) (GenBank accession no. WEY08940) are shown. The predicted signal sequences (amino acids 1-17) and transmembrane domains (amino acids 537-560) are depicted with a grey background. Amino acid positions differing between the two HA antigens are shown in bold letters. The amino acid identity of the two HA antigens is 95.5%.

**Supplementary Fig. 3.**
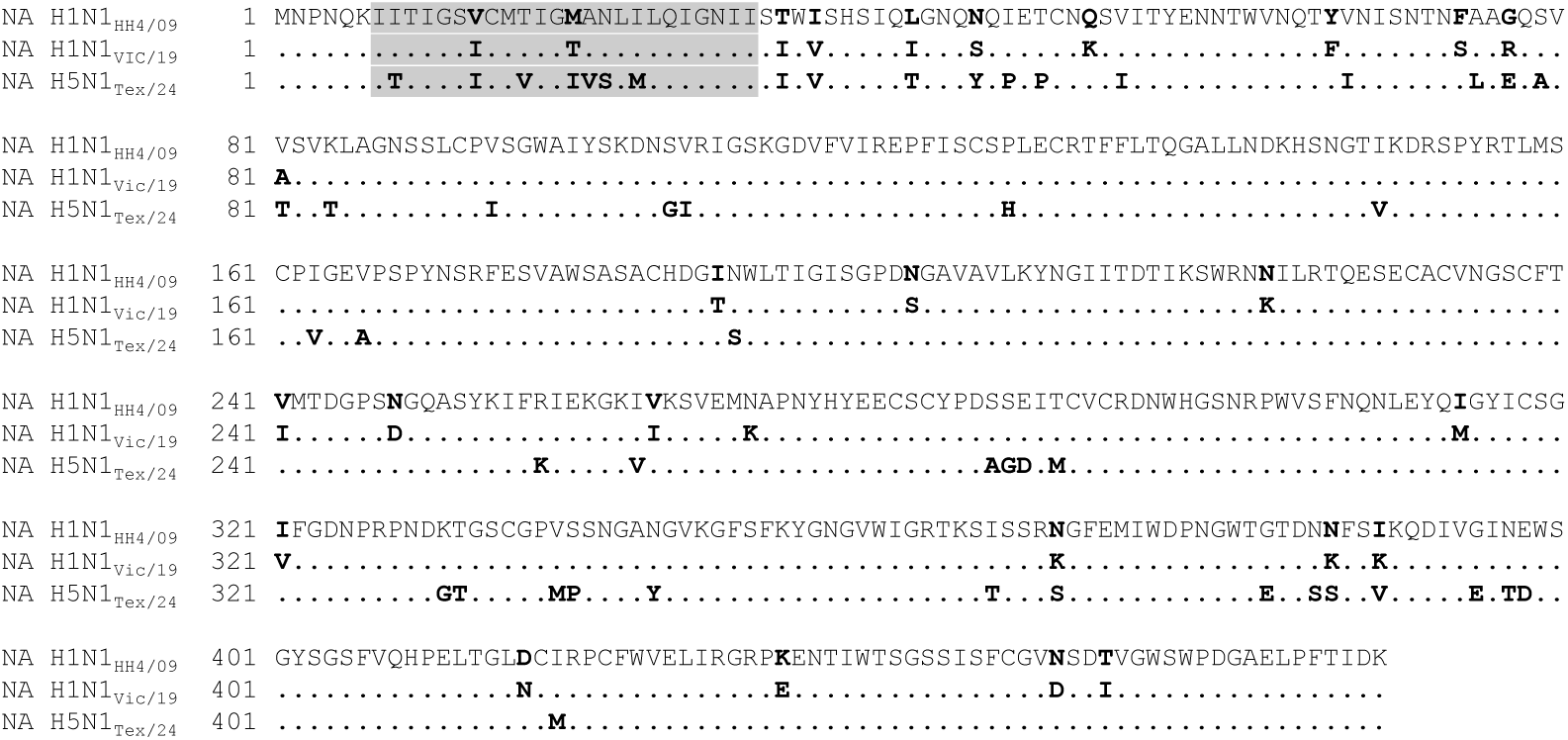
Alignment of NA primary amino acids sequences. Alignment of the NA amino acid sequences of A/Hamburg/4/2009 (H1N1) (GenBank accession no. ACR10227), A/Victoria/2570/2019 (H1N1) (GenBank accession no. WEY08939), and A/cattle/Texas/063224-24-1/2024 (H5N1) (GISAID acc. no. EPI_ISL_19155861) are shown. The predicted transmembrane domains (amino acids 7-30) are depicted with a grey background. Amino acid positions differing between the two NA antigens are shown in bold letters.

**Supplementary Table 1.** Clinical score of SPF pigs following boost vaccination.

| Animal no. | Clinical score at the indicated day of the experiment <sup>a</sup> |  |  |  |  |  |  |  |  |
| --- | --- | --- | --- | --- | --- | --- | --- | --- | --- |
|  | d28 | d29 | d30 | d31 | d32 | d33 | d34 | d35 | d36 |
| A1 | 0 | 2 | 1 | 0 | 0 | 0 | 0 |  |  |
| A2 | 0 | 2 | 1 | 0 | 0 | 0 | 0 |  |  |
| A3 | 0 | 1 | 1 | 0 | 0 | 0 | 0 |  |  |
| A4 | 0 | 2 | 1 | 0 | 0 | 0 | 0 |  |  |
| A5 | 0 | 2 | 1 | 0 | 0 | 0 | 0 |  |  |
| B1 | 0 | 2 | 1 | 0 | 0 | 0 | 0 |  |  |
| B2 | 0 | 0 | 1 | 0 | 0 | 0 | 0 |  |  |
| B3 | 0 | 0 | 1 | 0 | 0 | 0 | 0 |  |  |
| B4 | 0 | 0 | 0 | 0 | 0 | 0 | 0 |  |  |
| B5 | 0 | 0 | 0 | 0 | 0 | 0 | 0 |  |  |
| C1 | 0 | 0 | 0 | 0 | 3 | 3 | 1 | 0 | 0 |
| C2 | 0 | 0 | 0 | 0 | 0 | 3 | 1 | 0 | 0 |
| C3 | 0 | 0 | 0 | 0 | 0 | 3 | 1 | 0 | 0 |
| C4 | 0 | 0 | 0 | 0 | 0 | 3 | 1 | 0 | 0 |
| C5 | 0 | 0 | 0 | 2 | 3 | 3 | 1 | 0 | 0 |
| SC1 <sup>b</sup> |  |  | 0 | 0 | 2 | 3 | 0 | 0 | 0 |
| SC2 |  |  | 0 | 0 | 0 | 3 | 2 | 0 | 0 |
| SC3 |  |  | 0 | 0 | 2 | 3 | 0 | 0 | 0 |
| D1 | 0 | 0 | 0 | 0 | 0 | 0 | 0 | 0 | 0 |
| D2 | 0 | 0 | 0 | 0 | 0 | 0 | 0 | 0 | 0 |
| D3 | 0 | 0 | 0 | 0 | 0 | 0 | 0 | 0 | 0 |
| D4 | 0 | 0 | 0 | 0 | 0 | 0 | 0 | 0 | 0 |
| D5 | 0 | 0 | 0 | 0 | 0 | 0 | 0 | 0 | 0 |
| SD1 |  |  | 0 | 0 | 0 | 0 | 0 | 0 | 0 |
| SD2 |  |  | 0 | 0 | 0 | 0 | 0 | 0 | 0 |
| SD3 |  |  | 0 | 0 | 0 | 0 | 0 | 0 | 0 |

<sup>a</sup> Clinical symptoms of disease were rated 0 (no symptoms), 1 (mild symptoms), 2 (moderate symptoms), 3 (severe symptoms). The total score corresponds to the cumulative sum of all symptoms rated and was assigned for each animal at each day of the survey.
<sup>b</sup> SC, sentinel pigs co-housed with group C animals.
<sup>c</sup> SD, sentinel pigs co-housed with group D animals.

**Supplementary Table 2.** Clinical scoring of SPF pigs following challenge infection with the H1N1_HH4/09_/H1N1_Vic/19_ 6:2 reassortant virus.

| Animal no. | Clinical score at the indicated day post challenge <sup>a</sup> |  |  |  |  |  |
| --- | --- | --- | --- | --- | --- | --- |
|  | d56 | d57 | d58 | d59 | d60 | d61 |
| **A1** | 0 | 0 | 1 | 1 | 0 | 0 |
| **A2** | 0 | 0 | 1 | 0 | 0 | 0 |
| **A3** | 0 | 0 | 1 | 1 | 0 | 0 |
| **A4** | 0 | 0 | 1 | 1 | 1 | 0 |
| **A5** | 0 | 0 | 1 | 1 | 1 | 0 |
| **B1** | 0 | 0 | 1 | 0 | 0 | 0 |
| **B2** | 0 | 0 | 1 | 0 | 0 | 0 |
| **B3** | 0 | 0 | 0 | 0 | 0 | 0 |
| **B4** | 0 | 0 | 1 | 0 | 0 | 0 |
| **B5** | 0 | 0 | 1 | 0 | 0 | 0 |
| **C1** | 0 | 0 | 0 | 0 | 0 | 0 |
| **C2** | 0 | 0 | 0 | 0 | 0 | 0 |
| **C3** | 0 | 0 | 0 | 0 | 0 | 0 |
| **C4** | 0 | 0 | 0 | 0 | 0 | 0 |
| **C5** | 0 | 0 | 0 | 0 | 0 | 0 |
| **D1** | 0 | 0 | 0 | 0 | 0 | 0 |
| **D2** | 0 | 0 | 0 | 0 | 0 | 0 |
| **D3** | 0 | 0 | 0 | 0 | 0 | 0 |
| **D4** | 0 | 0 | 0 | 0 | 0 | 0 |
| **D5** | 0 | 0 | 0 | 0 | 0 | 0 |

<sup>a</sup> Clinical symptoms of disease were rated 0 (no symptoms), 1 (mild symptoms), 2 (moderate symptoms), 3 (severe symptoms). The total score corresponds to the cumulative sum of all symptoms rated and was assigned for each animal at all days after challenge infection.

## Notes

https://doi.org/10.5281/zenodo.17302034

### Competing Interest Statement

Robin Avanthay, Gert Zimmer, and Artur Summerfield filed a patent application related to the intramuscular prime/intranasal boost vaccine described in this work. All other authors declare no competing interests.

